# Genomic and Transcriptomic Analysis of Ninein Alternative Splicing Between C57BL/6J and DBA/2J Mice

**DOI:** 10.1101/2024.05.16.594557

**Authors:** ER Gnatowski, JL Jurmain, M Dozmorov, JT Wolstenholme, MF Miles

**Author notes:** Corresponding author: Michael Miles, MD, PhD, Professor, Department of Pharmacology and Toxicology Virginia Commonwealth University, **Email:**.

## Abstract

Ethanol’s anxiolytic actions contribute to increased consumption and the development of Alcohol Use Disorder (AUD). Our laboratory previously identified genetic loci contributing to the anxiolytic-like properties of ethanol in BXD recombinant inbred mice, derived from C57BL/6J (B6) and DBA/2J (D2) progenitor strains. That work identified Ninein (*Nin*) as a candidate gene underlying ethanol’s acute anxiolytic-like properties in BXD mice. *Nin* has a complex exonic content with known alternative splicing events that alter cellular distribution of the NIN protein. *We* hypothesize that strain-specific differences in *Nin* alternative splicing contribute to changes in *Nin* gene expression and B6/D2 strain differences in ethanol anxiolysis. Using quantitative reverse-transcriptase PCR to target *Nin* alternative splicing, we identified isoform-specific exon expression differences between B6 and D2 mice in prefrontal cortex, nucleus accumbens and amygdala. We extended this analysis using deep RNA sequencing in B6 and D2 nucleus accumbens samples and that *Nin* expression was significantly higher in D2 mice. Furthermore, exon utilization and alternative splicing analyses identified 8 differentially utilized exons and significant exon-skipping events between the strains, including 3 novel splicing events in the 3’ end of the *Nin* gene that were specific to the D2 strain. Our studies provide the first in-depth analysis of *Nin* alternative splicing in brain and identify a potential genetic mechanism altering *Nin* expression between B6 and D2 mice, thus contributing to differences in the anxiolytic-like properties of ethanol between these strains. This work contributes to our understanding of genetic differences modulating ethanol actions on anxiety that may contribute to the risk for alcohol use disorder.

## Introduction

The acute behavioral responses to ethanol in both humans and animal models include anxiolysis. Both clinical and preclinical research has demonstrated that the anxiolytic effects of acute ethanol consumption can lead to increased alcohol intake and play a critical role in the development of alcohol use disorder (AUD) [1; 2]. Ethanol’s anxiolytic properties are believed to enhance its reinforcing effects, resulting in a more pronounced response to ethanol in individuals who exhibit greater susceptibility to stress or anxiety compared to those with lower levels of anxiety [2; 3]. This enhanced anxiolytic response could result in a greater predisposition to consume alcohol, ultimately aiding development of alcohol use disorder (AUD). Furthermore, ethanol’s anxiolytic properties generate a potential feed-forward action on ethanol consumption due to the increase anxiety seen upon withdrawal in AUD subjects [3; 4]. The genetic variance underlying ethanol’s anxiolytic effects could thus influence both the development of AUD as well as relapse during ethanol withdrawal.

Ninein (*Nin*) was previously identified through a 2-stage quantitative trait loci (QTL) mapping in BXD recombinant inbred (RI) mice as a Chr 12 candidate gene underlying ethanol’s anxiolytic-like properties in the light-dark box transitional model of anxiety [5]. BXD RI strains are derived from C57BL/6J (B6) and DBA/2J (D2) inbred progenitor mouse strains which exhibit contrasting responses across multiple ethanol-related behaviors [6]. Confirmational analysis of *Nin* as a candidate gene revealed higher mRNA expression in D2 progenitors compared to B6, suggesting that higher *Nin* expression reduced the anxiolytic-like actions of ethanol in the light-dark box [5]. Protein expression analysis identified significantly higher expression of 2 provisional Ninein isoforms in D2 mice compared to B6 animals, thus implicating potential differences in splicing between these two strains [5]. However, the full documentation and characterization of these potential splicing differences in *Nin* expression between B6 and D2 mice remain to be determined.

Ninein is a microtubule-associated protein that plays a role in microtubule organization at the centrosome and in the cytoplasm [7; 8]. Ninein associates with the centrosome in many cell types, where it recaptures minus-ends of microtubules and is essential for apico-basal microtubule formation characteristic of complex polarized cells such as neurons [8; 9]. Ninein has primarily been studied in the context of development and has been demonstrated to undergo extensive alternative splicing that influences neural progenitor cells (NPCs) differentiation into neurons [10]. Previous published reports have identified roles of specific *Nin* exons that contribute localization and function during development. Understanding the role of *Nin* alternative splicing in adult animals may contribute to characterizing its role in regulating ethanol’s anxiolytic properties.

RNA splicing is a post-transcriptional modification that plays a critical role in mammalian gene expression [11]. This process involves the removal of introns from newly transcribed pre-mRNA sequences and is essential for generating mature protein-coding mRNAs. Compared to constitutive splicing, alternative splicing events comprise multiple processes that result in the inclusion or exclusion of specific exons in numerous combinations leading to the generation of multiple unique mRNA and protein isoforms from a single gene [12]. These processes generate transcriptome-wide complexity that can occur under both physiological and pathophysiological conditions. The human brain demonstrates a significantly higher occurrence of alternative splicing events that are evolutionarily conserved, demonstrating a functional role of alternative isoforms in the brain [13; 14]. There are five basic modes of alternative splicing: exon skipping (or cassette-type alternative exon), mutually exclusive exons, alternative 3’ splice site, alternative 5’ splice site, and intron retention. Exon skipping is the most prevalent pattern of alternative splicing in vertebrates and invertebrates [15].

Our prior work strongly implicates *Nin* as a candidate gene underlying a Chr 12 QTL for ethanol anxiolytic-like behavioral responses in BXD mice, and that this may involve both genetic regulation of Nin expression and splicing. In order to characterize potential mechanisms of *Nin* expression differences more fully between the BXD progenitor strains, B6 and D2 mice, here we studied strain-specific differences in exon utilization and splicing using selective quantitative RT-PCR and RNAseq analysis. By conducting deep sequencing of RNA transcripts, we identified novel strain-specific *Nin* alternative splicing and exon-utilization events in B6 and D2 adult mice. Our data demonstrate strain differences in previously reported exon 18 expression and splicing, as well as inclusion of a novel protein coding exons in the D2 strain within the 3’ region of the genome. Furthermore, we confirm potentially important coding region polymorphisms in D2 mice that might also alter *Nin* splicing and function. This evaluation of alternative splicing of a QTL derived candidate gene provides an initial framework for investigating the functional basis of strain-specific gene expression and demonstrates the potential complexity of genetic differences modulating complex behaviors such as AUD.

## Methods

### Animal Subjects

Male C57BL/6J and DBA/2J mice were obtained from the Jackson Laboratory (Bar Harbor, ME, USA) at 7-8 weeks of age and habituated to the vivarium for at least 1 week prior to initiating experimental studies. Animals were group-housed (four animals per group with ad libitum access to food and water, under a 12-hour light/dark cycle, in a 21 °C environment). Experiments were done in the light phase. All experiments were approved by the Institutional Animal Care and Use Committee of Virginia Commonwealth University and followed the National Institutes of Health Guidelines for the Care and Use of Laboratory Animals. At 9 weeks of age, animals were sacrificed by cervical dislocation and decapitation. Immediately thereafter, brains were extracted and chilled for 1 minute in cold phosphate buffered saline before brain regions being micro-dissected as previously described [16]. Excised regions were placed in individual tubes, flash-frozen in liquid nitrogen, and stored at −80 °C. Medial prefrontal cortex (mPFC), Nucleus Accumbens (NAc), and Amygdala (Amy) sections were used for RNA extraction and analysis as below.

### RNA extractions

Samples were homogenized with a Polytron® (Kinematica AG, Malters, Switzerland) and total RNA was extracted using a guanidine/phenol/chloroform method (STAT-60, Tel-Test, Inc. Friendswood, TX, United States) as per manufacturer guidelines. Each RNA liquid layer was added to a miRNeasy Mini Column (Cat #: 217004, Qiagen, Hilden, Germany) for cleanup and elution of total RNA. RNA concentration was determined by measuring absorbance at 260 nm using a Nanodrop 2000 (Thermo Scientific). RNA quality and purity was assessed by 260/280 absorbance ratios and by electrophoresis using the Agilent 2100 Bioanalyzer (Agilent Technologies, Savage, MD, United States). Samples all had RNA Integrity Numbers (RIN) ≥ 8 and 260/280 ratios between 1.96 and 2.05.

### Quantitative Reverse Transcriptase PCR

500 ng of RNA was converted to cDNA using the iScript cDNA synthesis kit (#1708891, Bio-Rad, Hercules, CA, United States) containing random hexamers according to the manufacturer’s instructions. All cDNA samples were diluted to 1ng/μL. qRT-PCR was performed using the Bio-Rad CFX Connect thermocycler according to manufacturer’s instructions for iTaq™ Universal SYBR® Green Supermix (#1725124, Bio-Rad, Hercules, CA, United States). Primers (Eurofin Genomics, Luxembourg) efficiencies were between 90-110% and each primer set resulted in only one PCR product on gel electrophoresis. Primer sequences, amplicon sizes, and Tm’s used for each gene are listed in **Supplemental Table 1**. Data analysis was performed using the 2-[ΔΔCT] method (Heid, 1996). Relative mRNA expression was normalized to housekeeping genes Ublcp1, Sort1, and Ppp2r2a, and Stab2 was used as a strain control. Ublcp1 was excluded in the amygdala after seeing significant strain differences in expression. Statistical analysis of qRT-PCR data was performed using a student’s t-test between strains within each brain region. For Gel Electrophoresis, PCR samples were combined with 1 μL of 6X gel loading dye (New England Biolabs, Cat. # B7024S) per 5 μL of sample and loaded each the remaining wells. 5 μL of TrackIt 50bp DNA ladder (Invitrogen, Cat. #10488043) was loaded into the first well. Electrophoresis was performed for 90 minutes with our power supply (BioRad PowerPac 300) set to 90V. 4% Agarose gels were run at room temperature (∼25°C). Gels were imaged using the Biorad GelDoc System (Bio-Rad, Cat. #1704486).

### RNAseq Library Preparation and Sequencing

RNAseq data have been deposited with the Gene Expression Omnibus resource (accession number pending). Library preparation and sequencing were completed by the VCU Genomics Core. A total of 5 biological replicates were obtained from each strain. Preparation of cDNA libraries was conducted using the standard protocols for the Illumina Stranded mRNA Prep, Ligation Kit (#20040534, Illumina, San Diego, CA, United States). Library insert size was determined using an Agilent Bioanalyzer. Libraries were sequenced on the Illumina NextSeq2000 (Illumina, San Diego, CA, United States) with 150 bp paired-end reads for a target depth of 100 million reads per sample. RNAseq data has been submitted to the Gene Expression Omnibus (GEO) database (accession number pending).

### RNAseq Quality Control and Alignment

Initial quality control checks were performed by FastQC (https://github.com/s-andrews/FastQC). All samples showed mean quality scores > 30. Fastp v. 0.22.0 [17] was used for adapter and end trimming and further quality control prior to alignment. B6 and D2 samples were aligned to release 108 of the Ensembl genome for C57BL/6J mice using STAR v 2.7.10b [18]. Aligned BAM files produced by STAR were further sorted by coordinate using Samtools v 1.6 [19]. Raw read counts for each BAM file were assigned and quantified using FeatureCounts (Samtools). A summary of RNAseq metrics can be found in Supplementary Table 2.

### Differential Gene Expression Analysis

Count files were analyzed for differential gene expression between B6 and D2 animals using the R package DESeq2 v 1.36.0 [20]. Low expressed genes where the median across all samples is zero were eliminated prior to analysis. Principal Component Analysis of the variance on the top 10,000 genes was run to identify significant sample outliers. Genes with a false discovery rate (FDR) < 0.05 were considered significantly altered and used in downstream analyses.

### Differential Exon Usage Analysis

A GFF annotation file containing collapsed exon counting bins was prepared from the UCSC GRCm39/mm39 GTF file using DEXSeq v. 1.42.10 [21]. Python script *dexseq_prepare_annotation.py*. The number of reads overlapping each exon bin was then counted using the DEXSeq Python script *dexseq_count.py*, the GFF file, and each sample’s BAM file. Differential exon usage (DEU) analysis was then carried out for the same contrasts studied in our DGE analysis using the DEXSeq R package standard analysis workflow. For analysis of Ninein differential exon usage, we utilized an FDR < 0.2 to identify significantly altered exon events.

### BXD Amygdala Exon-Level Expression Data

To further evaluate the relationship between Ninein exon expression and acute ethanol anxiolytic-like activity, we compared our DEXSeq results with the amygdala exon-level dataset in GeneNetwork (INIA Amygdala Exon Affy MoGene 1.0 S T (Nov10) RMA) [22]. This dataset provides estimates of mRNA expression in the Amygdala of 58 genetically diverse strains including 54 BXD Recombinant inbred strains, two F1 hybrid strains (B6D2F1 and D2B6F1), and the two progenitor strains (C57BL/6J and DBA/2J). 33 coding exons were correlated with the Top 500 BXD Phenotypes and filtered for Pearson correlations with previous ethanol light-dark box ethanol anxiolytic-like activity data from our prior studies in BXD mice [5]. Exons were correlated with acute ethanol phenotypes from the light dark box (LDB) transitional model of anxiety for percent distance traveled in the light (%DTL) or percent time spent in the light (%TIL) in response to an acute dose of ethanol (1.8 g/kg) at either 5-minute or 10-minute intervals.

### Alternative Splicing Analysis

Analysis of *Nin* exon utilization and splicing focused on five fundamental alternative splicing events: exon skipping, alternative 5′ and 3′ splice sites, mutually exclusive exons, and intron retention. Junction read counts for alternative splicing events were quantified by rMATS v. 4.1.2 [23] utilizing the STAR alignment BAM files as input. An FDR cutoff of 0.2 was used to identify significant alternative splicing events. P-values and FDRs that are smaller than the numerical accuracy cutoff (p > 2.2E-16) register as 0 in the rMATs output **(see Supplementary Table 4).** Rmats2shasimiplot v 2.0.4 (rMATs) and Maser v 1.14.0 [24] were used to analyze and visualize rMATs outputs.

Regtools v. 0.5.2 [25] was used to quantify junction reads and annotate novel/unusual junctions using the “junctions annotate” function. Outputs include chromosome junction start and end coordinates, as well as a score indicating the number of reads supporting the junction. Junctions were filtered again to identify novel splice junctions contain either a novel donor (D), a novel acceptor (A), a novel donor and novel acceptor pair (NDA), or no known donor or acceptor (N). Junctions with both a median score greater than 1 and a mean score greater than 1 across all 10 samples were included for statistical analysis. Scores across all 10 samples were compared using a student’s *t*-test and Bonferroni correction for multiple testing across the number of junctions passing filtering.

### Single Nucleotide Polymorphism (SNP) Analysis

To analyze single nucleotide polymorphisms (SNPs), we used the Mouse Genome Informatics (MGI) [26; 27] and Ensembl [28] dates using the GRCm39 assembly. We used the MGI database to output specific SNPs differing in the *Nin* gene from chr 12: 70,058,297-70,159,961. We used the Ensembl Database to identify the variant type and overlapping regulatory regions.

## Results

### D2 Mice Exhibit Increased Ninein mRNA Expression

We selected C57BL/6J (B6) and DBA/2J (D2) mice for our studies because of their distinct ethanol behaviors and status as progenitor strains for the BXD RI strains that were used in identification of *Nin* as a candidate gene for ethanol-anxiolysis. Total RNA was isolated from ethanol-naïve male B6 and D2 mice from the nucleus accumbens (NAc), medial prefrontal cortex (mPFC), and amygdala (AMY) for evaluating *Nin* alternative splicing events using qRT-PCR **(Figure 1).** Primers were designed to target different *Nin* exons reflective of known *Nin* transcript variants. Primer labels containing a (+) represent the inclusion of that exon in the primer design. Primer labels containing a (-) represent the exclusion of that exon from primer design and subsequent transcripts. These primer targets are defined according to the canonical Nin transcript (Fig. 2; Ensemble ID: ENSMUST00000085314.11) and included the canonical 3’ untranslated region, the alternative 5’ splice site at exon 28 (Ensemble IDs: ENSMUST00000085314.11 and ENSMUST00000222237.2), transcripts excluding exon 18 (Ensemble IDs: ENSMUST00000220689.2 and ENSMUST00000222835.2), and transcripts including exon 29 (Ensemble IDs: ENSMUST00000220689.2 and ENSMUST00000223257.2) (Figure 1, Supplementary Table 1). Primers investigating exon 28 targeted the shortened splice form of the exon.

**Figure 1.**
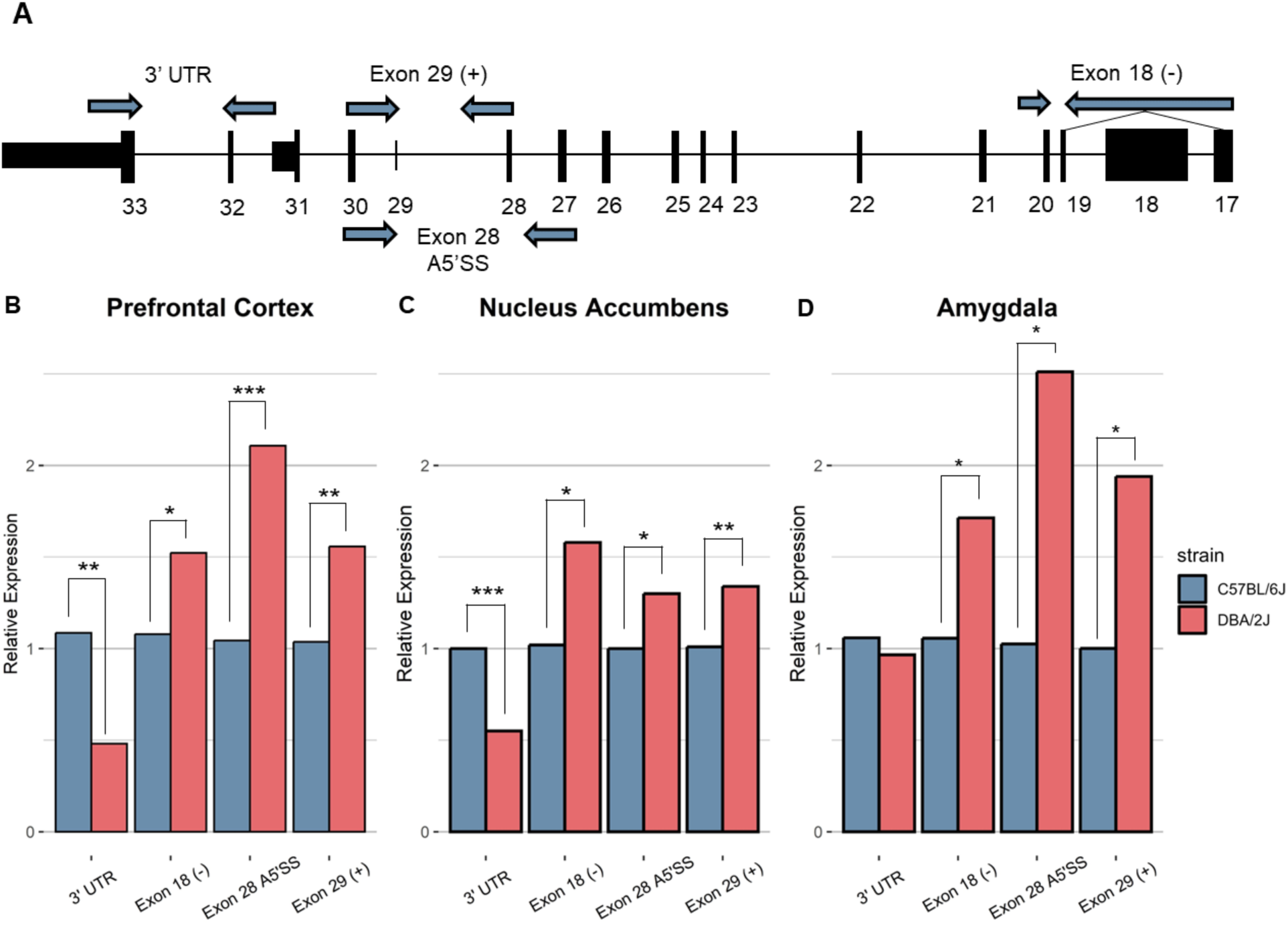
Ninein Exon-level qRT-PCR. Ninein exon-level mRNA expression between B6 and D2 was measured across the (B) prefrontal cortex, (C) nucleus accumbens, and (D) amygdala. (A) Representative image of *Nin* canonical transcript with primer positions. Basal mRNA levels for primers excluding Exon 18, including Exon 29, and using the alternative 5’ splice site for Exon 28 were significantly greater in D2 mice than B6 mice across all brain regions (* P < 0.01, ** P < 0.001, *** P < 0.0001, n = 5-10 per strain per brain region, Student’s *t*-test between strains). Primers amplifying the 3’ untranslated region (UTR) showed significantly higher expression in B6 mice compared to D2 mice in the PFC and NAc.

**Figure 2.**
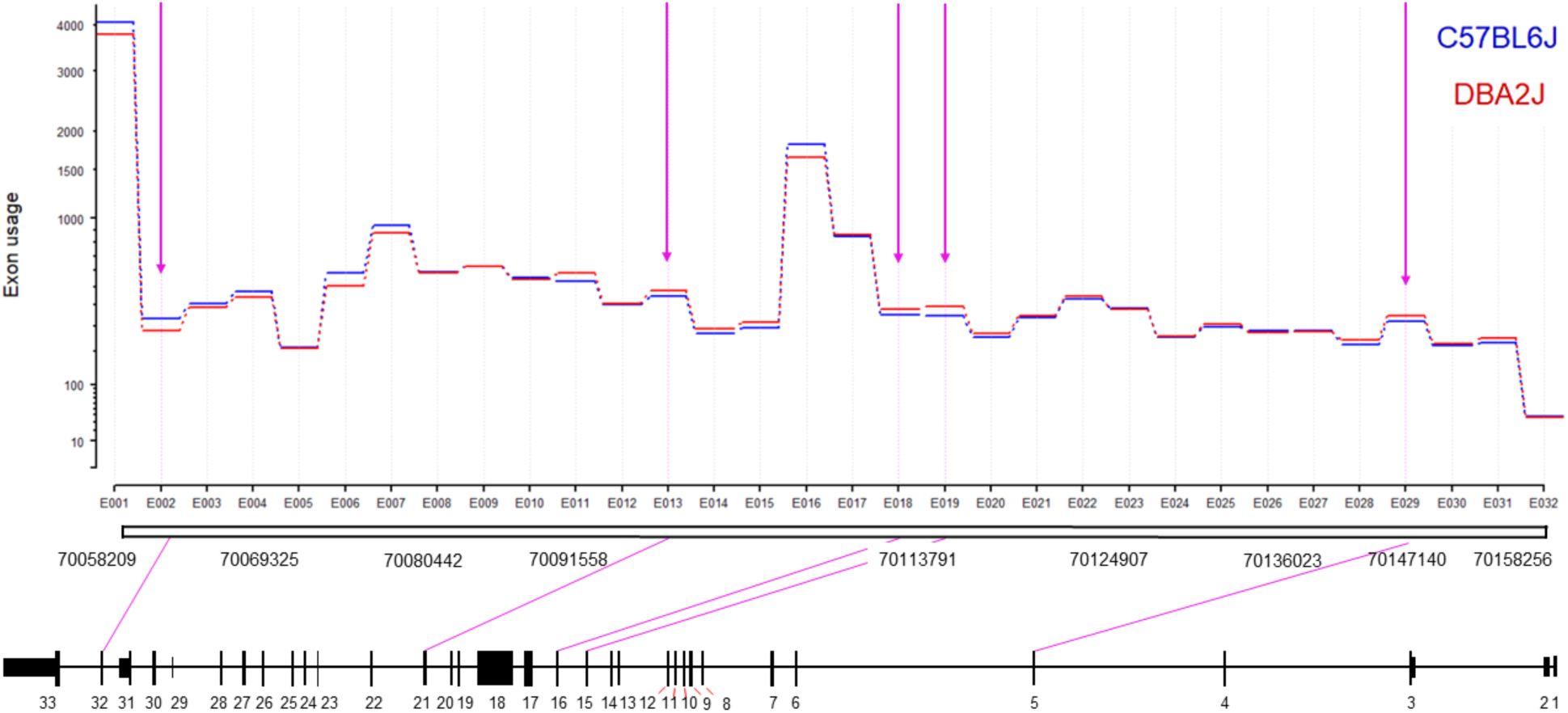
Ninein Differential Exon Utilization. The y-axis indicates level of exon usage on a logarhtmic scale. We identified 8 significant DEU events between B6 and D2 mice (FDR = 0.2). D2 mice showed significantly higher utilization of exons 5, 15, 16, 19, and 21. B6 mice showed significantly higher utilization of exon 32, and showed higher usage of retained introns between exons 9 and 10 and between exons 25 and 26.

In mPFC **(Figure 1A)**, D2 mice (n = 10) exhibited a significantly higher expression for exon 18 (-) (p = 0.021), exon 28 A5’SS (p = 2.85E-05) and exon 29 (+) (p = 0.0047). B6 mice (n = 10) showed significantly higher expression of the 3’ UTR region (p = 0.0038) of *Nin*. This same pattern of expression was seen in the NAc **(Figure 1B)** where D2 mice (n = 5/strain) showed higher expression of exons 28 A5’SS (p = 0.01), exon 29 (+) (p = 0.009), and exon 18 (-) (p = 0.02), and B6 mice showed significant higher expression of the 3’ UTR region (p = 0.00011). In the Amygdala **(Figure 1C)**, D2 mice (n = 5) showed the same pattern of expression for exon 28 A5’SS (p = 0.03), exon 29 (+) (p=0.033), and exon 18 (-) (p = 0.046) compared to B6 mice (n = 5), but there was no significant strain difference for the 3’ UTR (p = 0.718) in Amygdala. This suggests D2 mice have significantly higher total *Nin* expression compared to B6 mice across most exons, but the decreases in 3’UTR expression in D2 mice predicts an alternative 3’UTR and greater expression of shortened transcripts in that strain compared to B6 mice.

### Ninein Exhibits Differential Exon Utilization

To extend the PCR analysis and possibly detect novel transcript variants, we performed deep-read RNA sequencing of the NAc to further evaluate alternative splicing differences between B6 and D2 mice. Differential gene expression analysis DESeq2 confirmed our a priori hypothesis and qRT-PCR data above that D2 mice exhibit significantly higher *Nin* expression compared to B6 (LFC = 0.271, padj = 5.50685E-07). Full analysis of these D2 v. B6 deep sequencing results will be reported elsewhere (Gnatowski and Miles, in preparation). We then performed an exon-level analysis using DEXSeq to better characterize exon-level expression differences detected by the qRT-PCR **(Figure 2, Supplementary Table 3)**. We used a relaxed statistical threshold (FDR ≤ 0.2) without LFC filtering to define differential exon utilization (DEU) for *Nin.* DEXSeq identified 5 differentially utilized exons between B6 and D2 mice. Of these 5 significant DEU events, D2 mice had increased usage of exon 5 (p_adj_ = 0.175, LFC = 0.11), exon 15 (p_adj_ = 0.0026, LFC=0.18), exon 16 (p_adj_ = 0.153, LFC = 0.105), and exon 21 (p_adj_ = 0.179, LFC = 0.1), while B6 showed significantly increased usage of exon 32 (p_adj_ = 0.1104, LFC = −0.245). The increased expression of exon 32 in the B6 mice is consistent with those seen in the qRT-PCR characterization of the transcripts with the full 3’ UTR (**Figure 1**).

### GeneNetwork Correlations of Ninein Exons and Ethanol-Anxiolysis Behaviors

In order to interrogate the relationship between exon-level expression and behavior, we used the GeneNetwork resource to examine correlations in BXD mice between Ninein exon-level expression and our prior acute ethanol anxiolysis behavior **(Table 1)**, given the extensive amount of BXD brain region expression and phenotypic data available at that site. Fortuitously, given their role in stress and anxiety, exon-level expression data for amygdala and hypothalamus were available for BXD strains. For %TIL in response to ethanol in the amygdala, we identified significant Pearson correlations with exon 4 through 7 and exon 17 **(Table 1).** For %TIL in response to ethanol in the hypothalamus, we identified significant Pearson correlations with exon 4 through 6 and exon 17. We did not see significant correlations of exon 7 in the hypothalamus with %TIL.

**Table 1.** GeneNetwork *Nin* Exon Phenotype Correlations. *Nin* exon expression was correlated against the top 500 BXD phenotypes from GeneNetwork (Pearson Correlation). Exons showing significant correlation with acute ethanol anxiolysis in the Light Dark Box were identified as exons of interest. Record ID refers to GeneNetwork expression data file number.

Figures/Tables
|  |  | Amygdala |  |  |  | Hypothalamus |  |  |  |
| --- | --- | --- | --- | --- | --- | --- | --- | --- | --- |
|  |  | %DTL (EtOH)<br>5-minute |  | %TIL (EtOH)<br>5 minute |  | %DTL (EtOH)<br>5-minute |  | %TIL (EtOH)<br>5 minute |  |
| Record ID | Exon | Corr. | P-Value | Corr. | P-Value | Corr. | P-Value | Corr. | P-Value |
| 10400838 | Exon 4 | N/A | N/A | -0.601 | 1.37E-02 | -0.695 | 9.00E-04 | -0.7506 | 3.46E-03 |
| 10400837 | Exon 5 | -0.577 | 4.96E-03 | -0.578 | 1.91E-02 | -0.7367 | 2.60E-04 | -0.7826 | 1.60E-03 |
| 10400836 | Exon 6 | -0.607 | 2.72E-03 | -0.601 | 1.39E-02 | -0.7453 | 1.90E-04 | -0.8738 | 5.19E-05 |
| 10400835 | Exon 7 | -0.637 | 1.44E-03 | -0.607 | 1.26E-02 | N/A | N/A | N/A | N/A |
| 10400825 | Exon 17 | 0.623 | 5.23E-04 | 0.767 | 1.96E-03 | 0.611 | 5.92E-03 | 0.672 | 1.46E-02 |

For %DTL in response to ethanol in the amygdala, we identified significant Pearson with exons 5 through 7 and exon 17. Exon 4 showed no significant correlations for %DTL **(Table 1)**. For %DTL in response to ethanol in the hypothalamus, we identified significant Pearson with exons 4 through 6 and exon 17. Exon 7 showed no significant correlations for %DTL **(Table 1)**. The microarray probe set for Exon 8 contained SNPs in D2 mice that could alter expression and thus this exon data was excluded. These results suggest that the expression of a cluster of *Nin* exons are responsible for the regulation of ethanol anxiolysis in the LDB.

### Ninein Exhibits Strain-Specific Alternative Splicing

Since Ninein has 18 predicted transcript variants, we used rMATs to analyze strain-specific differences in alternative splicing. An FDR cutoff of ≤ 0.2 was employed along with a percent spliced in (PSI, ψ) cutoff of ψ > 0.05. rMATs analysis revealed 7 significant alternative splicing events between B6 and D2 mice, including 4 exon skipping events and 3 mutually exclusive exons **(Figure 3, Supplementary Tables 4-5).** For the exon skipping events, we identified a significant change in ψ values for the largest exon, exon 18 (FDR = 0.177, ψ = 0.066), and identified 3 novel exon skipping events only present in D2 mice occurring between exons 32 and 33 **(Figure 3A).** These exons will be referred to as 32A (FDR < 2.2e-16, Δψ = −0.153), 32A’ (FDR < 2.2e-16, Δψ = −0.164), and 32B (FDR < 2.2e-16, Δψ = −0.061). Exon 32A’ represents the alternative 5’ splicing of the larger identified exon, exon 32A. These identified exon skipping events overlap with the identification of transcripts where two exons were deemed mutually exclusive (MXE). Two of these mutually exclusive exon events identified exon 32B to be mutually exclusive from both 32A (FDR = 0.0051) and 32A’ (FDR = 0.00092) **(Figure 3B).** The third alternative splicing exon event showed a significant greater mutually exclusivity between exons 18 and 19 (FDR = 0.0605) in the B6 mice (**Figure 3B**).

**Figure 3.**
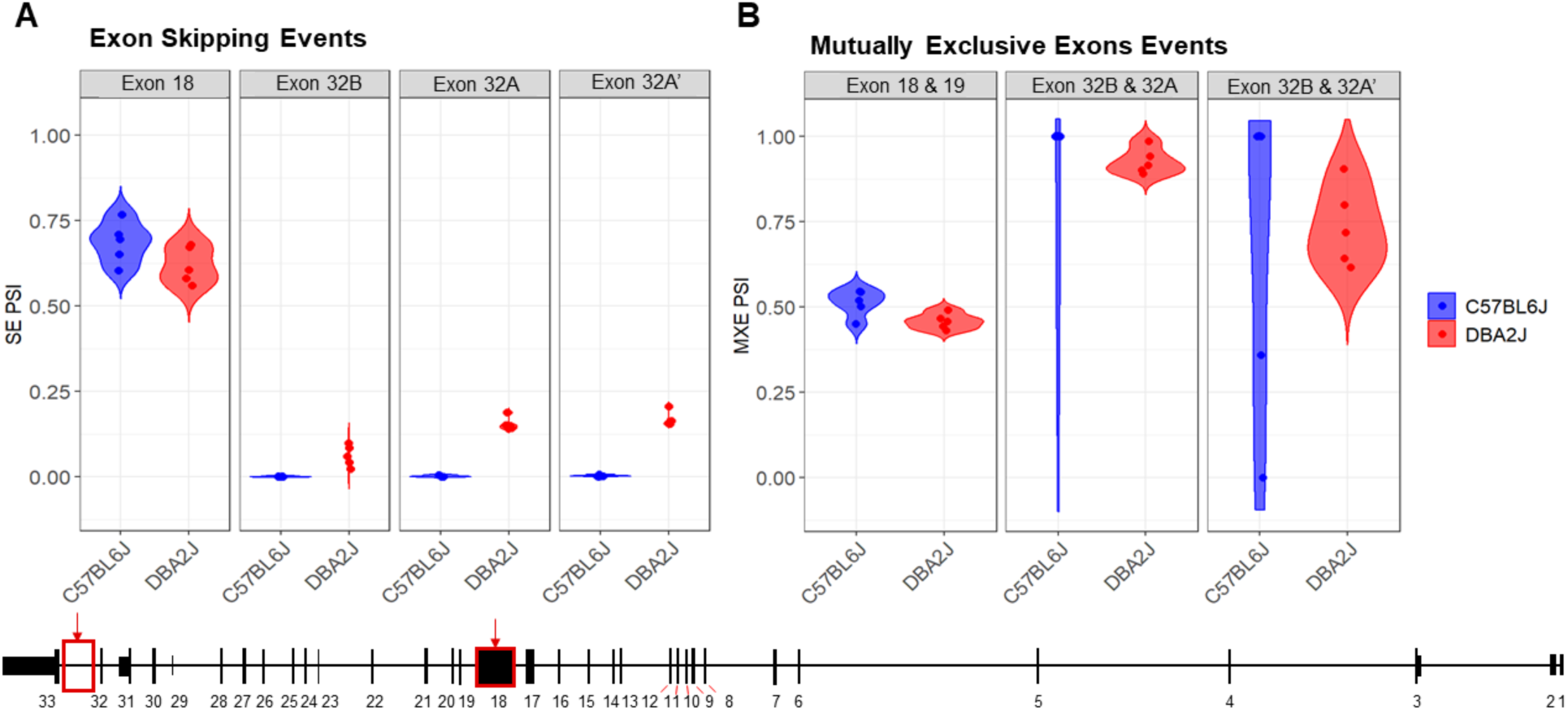
Ninein Alternative Splicing Events. Comparison of B6 and D2 *Nin* Alternative splicing events resulted in the identification of (A) 4 exon skipping events and (B) 3 mutually exclusive exon events (FDR ≤ 0.20, deltaPSI ≤ 0.05). Alternative splicing analysis identified 3 novel exon events exclusive to the DBA/2J strain (Ex 32B, PSI = −0.061, FDR = 0; Ex 32A, PSI = −0.153, FDR = 0; Ex 32A’, PSI = −0.164, FDR = 0).

To further investigate these alternative splicing events, we used *regtools* to characterize and quantify reads at novel junctions contributing to novel splicing events. We identified 2 significant splicing junctions containing either a novel splice acceptor or novel splice donor between exons 32 and 33 **(Figure 4**, **Table 2)**. The first significant junction identified contains a known splice acceptor on exon 33 (chr12:70,063,321) that spans to the 3’ end of identified novel exon 32B (chr12:70,064,347), which contains the novel donor. The second significant junction consists of a novel acceptor site on identified novel exon 32A/A’ (chr12:70,061,730) with a known splice donor on Exon 32 (chr12:70,062,498). These findings confirm that the splicing of known coding exons (32 and 33) is occurring at intronic sites specifically in D2 mice.

**Figure 4.**
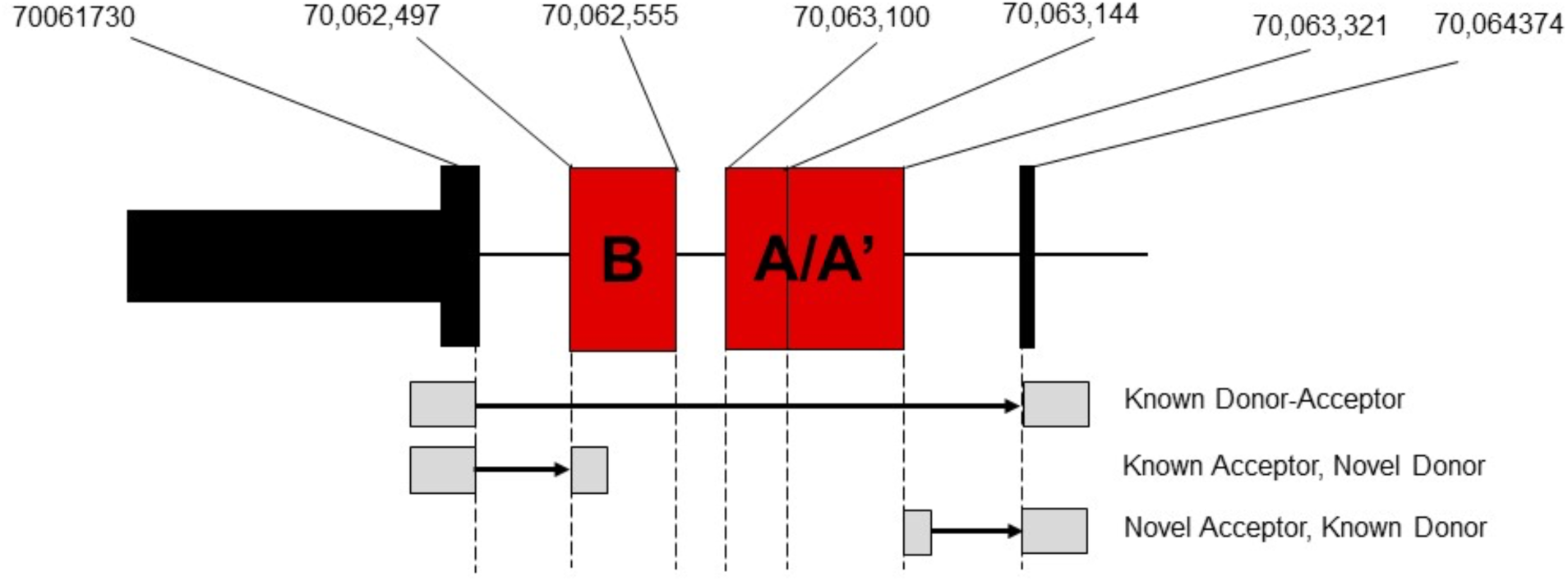
Junction Analysis of Novel *Nin* exons. Location of novel exons identified by the rMATs alternative splicing analysis and corresponding junctions identified by regtools.

**Table 2.** Junction Analysis of Novel *Nin* exons.

| **Start** | **End** | **Stran**<br>**d** | **Ancho**<br>**r** | **Average**<br>**B6 Score** | **Average**<br>**D2 Score** | **P-Value** | **Padj.** |
| --- | --- | --- | --- | --- | --- | --- | --- |
| 7006332<br>1 | 7006434<br>7 | - | D | 1.8 | 77 | 0.0005 | 0.003 |
| 7006173<br>0 | 7006249<br>8 | - | A | 0.4 | 22.6 | 0.003 | 0.019 |
| 7010804<br>9 | 7010946<br>3 | - | D | 2.2 | 6 | 0.033 | 0.196 |
| 7010950<br>2 | 7013731<br>0 | - | NDA | 2.2 | 8 | 0.134 | 0.805 |
| 7006173<br>0 | 7006310<br>1 | - | A | 0.2 | 3.4 | 0.011 | 0.066 |

**Table 3.** Nin Exon 18 Ensembl SNPs Between B6 and D2 strains.

| SNP ID | Coordinate | Allele Summary | B6 | D2 | Variant Type |  |
| --- | --- | --- | --- | --- | --- | --- |
| rs32227977 | 70089171 | C/T |  | C | Missense Variant |  |
| rs32227980 | 70089195 | G/T |  | T | Missense Variant |  |
| rs32227983 | 70089251 | A/G | A | G | Missense Variant | * |
| rs32228806 | 70089348 | A/G |  | G | Missense Variant |  |
| rs29183409 | 70089601 | A/G | A | G | Synonymous Variant |  |
| rs32224596 | 70089643 | A/T |  | T | Synonymous Variant |  |
| rs32224599 | 70089718 | C/T |  | T | Synonymous Variant |  |
| rs32224602 | 70089817 | C/T |  | T | Synonymous Variant |  |
| rs32225355 | 70089904 | C/T |  | C | Synonymous Variant |  |
| rs32225358 | 70089951 | C/T |  | T | Missense Variant |  |
| rs29202173 | 70089964 | A/G |  | A | Synonymous Variant |  |
| rs29192398 | 70090160 | C/T | C | T | Missense Variant | * |
| rs29159683 | 70090163 | G/T | G | T | Missense Variant |  |
| rs32225363 | 70090252 | C/T |  | T | Synonymous Variant |  |
| rs32226356 | 70090387 | A/G |  | G | Synonymous Variant |  |
| rs32226359 | 70090395 | C/T |  | C | Missense Variant |  |
| rs32226362 | 70090398 | C/T |  | T | Missense Variant |  |
| rs32227145 | 70090555 | C/T |  | C | Synonymous Variant |  |
| rs29149025 | 70090689 | C/T |  | C | Missense Variant |  |
| Rs32227150 | 70090905 | A/G |  | G | Missense Variant | * |

The identification of these novel splice junctions characterized two independent novel exons, labeled exons 32A and 32B **(Figure 5A),** with one of these novel exons undergoing alternative 5’ splicing (exon 32A’). We used the Integrative Genome Viewer (IGV) tool to analyze for possible open reading frames of the novel exons identified in the alternative splicing analyses. **Figure 5A** shows representative B6 and D2 samples to demonstrate the increased sequencing read density between exons 32 and 33 in D2 samples compared to B6 samples. IGV showed an exon present between exons 32 and 33 that aligns with the chromosomal location of the novel exon 32B (chr12:70,062,497-70,062,555) identified by rMATs. This exon contains a short reading frame of 5 amino acids followed by a termination codon and an alternative 3’ UTR (**Figure 5B**). For exon 32A and its alternative 5’ spliced counterpart, exon32A’, all theoretical reading frames contained multiple premature termination codons (**Figure 5C**). The longest possible reading frame had 46 amino acids prior to a termination codon at chr12:70,063,184. IGV also identified a guanine to adenine (G-A) SNP in the exon 32A region at chr12:70,063,271, however this SNP is a synonymous variant that maintains the status of the potential reading frames.

**Figure 5.**
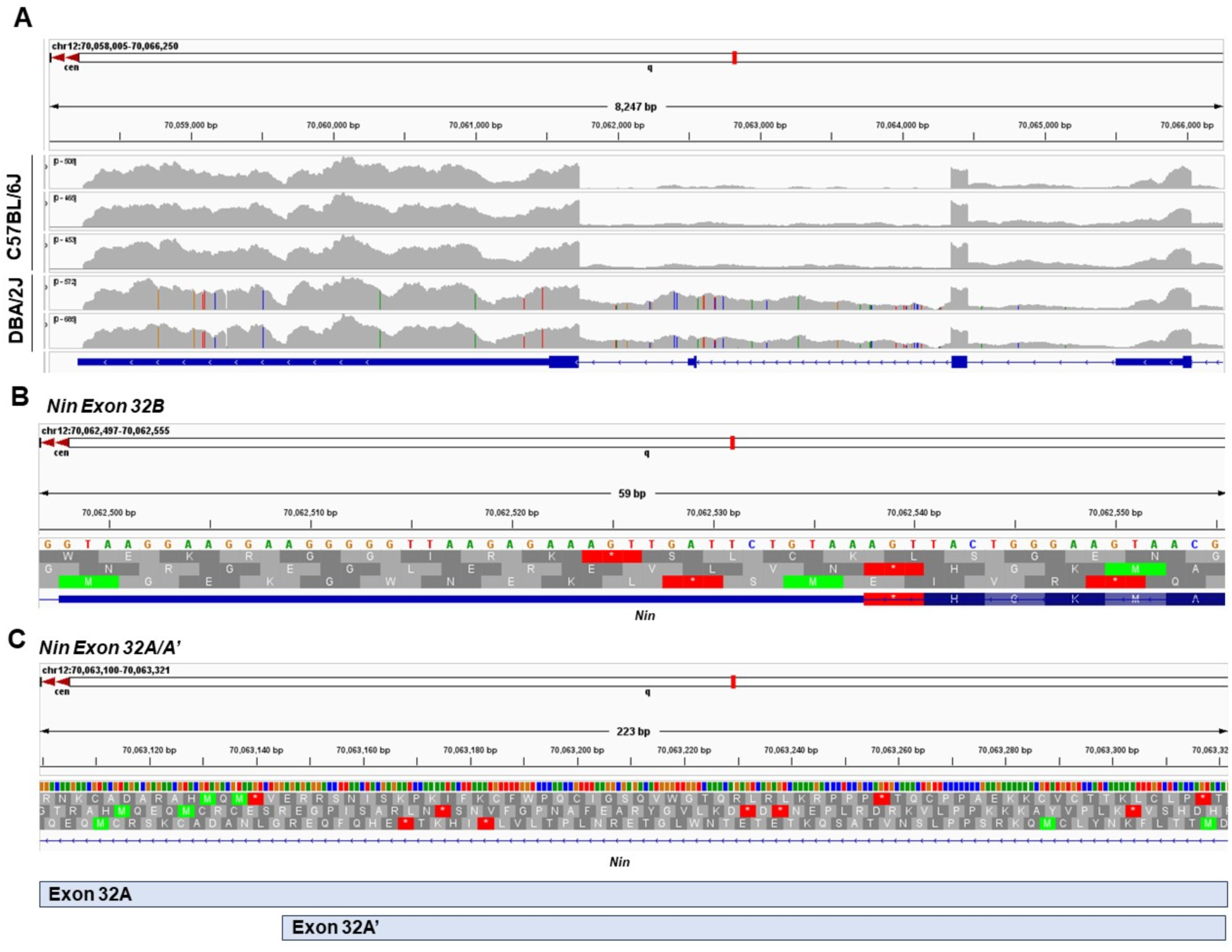
Intergrative Genome Viewer (IGV) Analysis of Novel Exon Open Reading Frame. **(A)** Comparison of Read distribution by strain for *Nin* exons 31 through 33. **(B)** Location of rMATs-identified novel exon 32B (chr12:70,062,497-70,062,555) including amino acid reading frames. **(C)** Location of rMATs-identified novel exons 32A (chr12:70,063,100-70,063,321) and 32A’ (chr12:70,063,144-70,063,321) including amino acid reading frames.

### PCR Validation of Novel Exons

In order to validate the supporting bioinformatic evidence for the existence of novel exons between exons 32 and 33, we performed RT-PCR targeted at these exons in D2 and B6 nucleus accumbens **(Figure 6).** PCR primers did not span exon-exon junctions to allow for the amplification between exons 32 and 33 **(Figure 6A)**. cDNA from representative B6 (n =3) and D2 (n =3) were size-separated by electrophoresis on 4% agarose. PCR products from D2 mice yielded two distinct bands: one band served as the representative band for the canonical splicing of exon 32 to exon 33 (214 bp) and a second larger band (∼250-300 bp) was identified, consistent with the predicted size of novel exon 32B **(Figure 6B)**. This additional band was not seen in the B6 samples. These results strongly support the existence of an exon exclusive to D2, located in the 3’ region of the gene.

**Figure 6.**
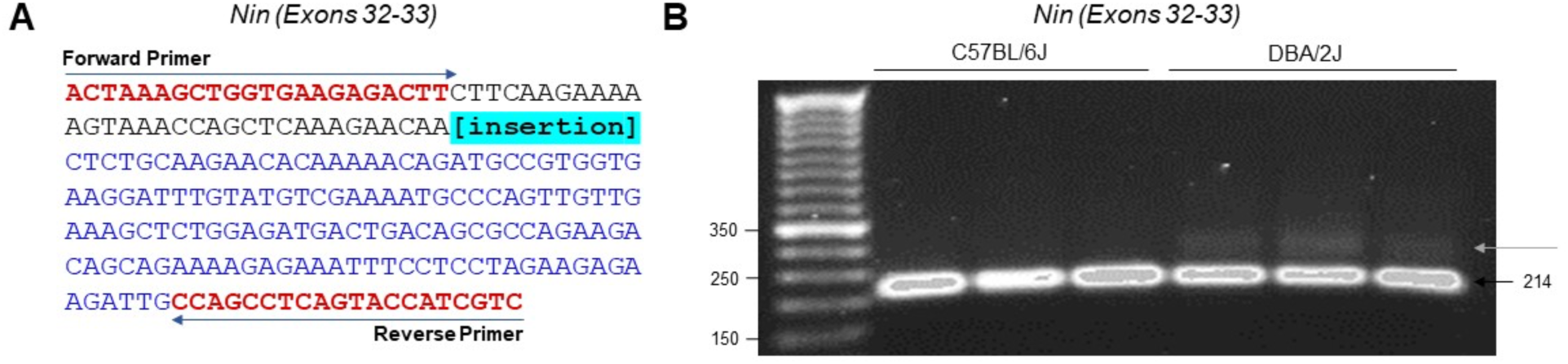
*Nin* Novel Exon Amplification via PCR and Gel Electrophoresis. **(A)** Sequence of the canonical PCR amplified region encompassing Exons 32 and 33. Amplicons of 214 and an unknown amplicon are obtained. The location of the novel insertion is highlighed in blue. The positions of the two primers (Forward and Reverse) were used for PCR amplification are shown (see Supplementary Table 1). **(B)** The target region of *Nin* was PCR amplifed using the previously described primers. PCR products were size-separated by electrophersis on a 4% agarose gel for 90-minutes.

### Ninein Single Nucleotide Polymorphisms

We identified 858 SNPs differing between the B6 and D2 strains in the *Nin* gene (chr 12: 70,058,297-70,159,961) using the MGI and Ensembl databases (GRCm39) **(see Supplementary Table 6)**. Our analysis identified 51 coding region SNPs, with 15 of these identified as missense variants. 4 of the missense variants were predicted to have “deleterious” effects on the amino acid sequence of the protein based on the SIFT score. 3 of these deleterious SNPs occur in exon 18 while the other occurs in exon 27. The other 11 missense SNPs were deemed “tolerated” by the SIFT algorithm, indicating the variant would not greatly impact protein function. 8 of these “tolerated” missense variants also occur within alternatively spliced exon 18. Variants *rs32226359*, *rs32226362*, and *rs29149025* occur in a known enhancer region (ENSMUSR00000789018). We also utilized this database to identify intronic SNPs that correspond to the regions associated with the proposed novel exons in D2 mice. There are 21 Intronic SNPs in the region of the proposed novel exons (chr12: 70,061,987-70,063,271) identified in the rMATs alternative splicing analysis. This region corresponds with a known enhancer region (ENSMUSR00000789014) in that area of the gene.

## Discussion

The studies presented here provide the first in-depth analysis of genetic variation in alternative splicing of *Nin*, a candidate gene for ethanol anxiolytic-like behavior. Deep RNA sequencing provided sufficient read depth to characterize strain-differences in exon utilization and alternative splicing events, allowing for the identificaion of novel strain-specific splicing events. PCR results generally indicated increased *Nin* expression in D2 mice, but a 50% reduction in *Nin* exon expression in in the 3’ UTR region of the gene. RNA sequencing both confirmed the increased usage of exon 32 in B6 mice and identified novel splicing events between exons 32 and 33. These novel splicing events indicated here pose the possibility of the use of alternative 3’ untranslated regions by the D2 mice which would result in truncated *Nin* protein isoforms. Our findings also showed increased exon skipping of the largest exon, exon 18, in D2 mice, which may occur as a consequence of the D2 strain containing 3 deleterious SNPs in this coding exon. The alternative splicing events characterized here may provide a genetic mechanism for strain specific regulation of *Nin* transcription contributing to behavioral differences in the anxiolytic response to an acute dose of ethanol between B6 and D2 mice. These results also provide further support for *Nin* being a candidate gene underlying the Etanq1 QTL previously identified by our laboratory for ethanol anxiolytic-like activity in BXD mice [5].

### Exon-Level Expression and Behavioral Correlations

Alternative splicing is a crucial step of post-transcriptional gene expression that substantially increases transcriptome diversity and is critical for diverse cellular processes, including cell differentiation, development, cellular localization, and tissue remodeling [22; 23]. Differential exon usage comes as a result of increased exon skipping, the most prevalent form of alternative splicing, leading to the selective inclusion or exclusion of coding regions during the formation of mRNA [11]. It has been proposed that RNA alternative splicing, specifically exon skipping, plays a causal role in Alcohol Use Disorder (AUD) susceptibility [14]. Our DEXSeq Analysis identified 5 differential exon usage events where 1 of these events overlapped with exon 5, which showed significant correlations with phenotypes from the anxiolytic response to ethanol previously identified in Putman et. al [5]. Exon 5 showed significant negative correlations with %TIL and %DTL in the light dark box in both the amygdala and hypothalamus, indicating that higher *Nin* exon 5 expression would result in a decreased anxiolytic response to ethanol. This correlation appears to pool in a strain-dependent manner where BXD strains with the B6 allele have decreased *Nin* expression **(see Supplementary Figure 1).** This is consistent with DEU results showing B6 mice have lower exon 5 usage compared to D2 mice. This highlights exons 5 as an exon-specific target underlying an ethanol-responsive candidate gene.

### Exon 18 Skipping

Ninein (*Nin*) is an exceptionally large gene containing 33 protein coding exons, generating protein products that can vary in an isoform-dependent manner. Previous work from Zhang and colleagues elucidated the role of alternative splicing of the *Nin* gene in the differentiation of neural progenitor cells (NPCs) into mature neurons [10]. They pinpointed two exons that play pivotal roles in this process. The first, a 61-nucleotide exon (exon 29), is specifically expressed in neurons but excluded in NPCs. The inclusion of this exon triggered the dissociation of *Nin* from the centrosomal protein CEP250, leading to *Ni* diffusion into the cytoplasm. Conversely, those investigators found that the exclusion of a larger exon (exon 18, >2,000 nucleotides) in neurons, that is not present in NPCs that resulted in the dissociation of another centrosomal protein, CEP170, from *Nin* [10]. Zhang and colleagues argued that these alternative splicing appeared sufficient to differentiate NPCs into neurons. Our initial characterization of *Nin* alternative splicing using qRT-PCR showed that D2 mice have increased expression of transcripts containing exon 29 and transcripts excluding exon 18 compared to B6. This suggests that D2 mice have higher expression of non-centrosomal *Nin* isoforms. This initial result was corroborated by the alternative splicing analysis that indicated exon 18 exhibited significantly greater exon skipping in D2 mice.

The decreased utilization of exon 18 in D2 mice may ultimately contribute to decreased localization of CEP170 to the centrosome and increased NPC differentiation into mature neurons. The underlying mechanism for D2 mice having a lower rate of exon 18 utilization is unknown. However, SNP analysis identified 11 missense variant SNPs in exon 18 alone, 3 of which were predicted “deleterious” SNPs that would result in a change in the protein structure leading to loss of function or harmful gain of function [29]. This could drive increased splicing out of exon 18 in D2 mice. However, even for *Nin* isoforms containing exon 18 in D2 mice, those proteins might not be fully functional in terms of centrosomal localization. The net result of decreased exon 18 utilization and probable decreased exon 18 function in D2 mice seems likely to impair *Nin* centrosome localization or function in non-neuronal cells and might lead to developmental alterations in the relative abundance of neurons in D2 mice given the role on *Nin* in neuronal differentiation. However, further confirmation of such differences between B6 and D2 mice at a cellular level are needed.

### Identification of Novel Exons and Alternative 3’ UTRs

Another major finding in from these studies is the identification of novel exons between exons 32 and 33 specifically expressed only in D2 mice. Our original findings from the qRT-PCR results looking at *Nin* exon expression highlighted a 50% reduction in the expression of the exon 32 to the 3’ UTR in D2 mice compared to B6 mice. This result was corroborated by the DEU results indicating that exon 32 has higher exon usage in B6 mice. Confirmation of the presence of these novel exons by PCR gel electrophoresis identified a second band exclusive to D2 mice consistent in size with inclusion of novel exon 32B, which contains an alternative stop codon and 3’ UTR. Together, these results indicate that D2 mice uniquely produce shortened *Nin* mRNA transcripts with alternate 3’ UTRs and and altered carboxy terminus of the protein coding region. It remains unclear whether or not the identification of novel exons 32A and 32A’ result in the inclusion of an additional 3’ UTR, or whether or not the inclusion of these exons leads to the increased utilization of a 3’UTR region following exon 31 shown in **Figure 5A**. This would be consistent with the lack of an additional band in the gel electrophoresis 177-221 bp higher than the known band.

### 3’ UTR- Localization

The 3’ untranslated region (3’ UTR) plays a pivotal role in post-transcriptional modulation of gene expression. The 3’ UTR often contains regulatory regions that influence gene expression by regulating processes such as polyadenylation, translation efficiency, localization, and stability of the mRNA [30; 31]. Given the *Nin* alternative exon usage in D2 mice noted above, it is possible that the alternative 3’ UTRs could change *Nin* mRNA localization or function. Transcripts containing multiple 3’ UTRs encoding the same protein have been repeatedly reported. For example, mRNAs for *BDNF* containing the same coding sequence with distance 3’ UTRs resulted in distinct differences in localization where the Long 3’ UTR localized BDNF to distal dendrites and shorter 3’ UTR *BDNF* transcripts remained in the soma [21]. Similar examples have been reported for *CamK2a* [32]. Of note, our prior studies on mRNA localization to synaptoneurosomes indicated increased *Nin* mRNA abundance in the cellular fraction containing pre- or post-synaptic contents in D2 mice [33]. In the case of *Nin*, the usage of alternative 3’ UTRs may provide a genetic mechanism for the regulating *Nin* trafficking and function in dendritic microtubule polarity, however the exact transcript altering this localization is unclear [31]. The localization of microtubules in dendrites may also contribute to changes in the trafficking of GABA and glycine receptors to the postsynaptic site [21; 34]. This highlights a specific cellular signaling mechanism by which *Nin* expression and splicing could modulate ethanol’s anxiolytic properties.

## Conclusion

This study is the first to provide an in-depth genomic and bioinformatic analysis of a QTL-identified candidate gene for ethanol anxiolytic activity. We observed strain-specific differences in exon regulation across different analysis parameters that we hypothesize played a critical role in regulating ethanol anxiolysis that lead to the identification of *Nin* as a candidate gene. The alternative splicing events that we identified could alter localization and expression of the NIN protein. *In vivo* studies examining deletion of specific *Nin* exons are needed to further validate the contribution of specifics exons in regulating *Nin* localization and expression and, ultimately, in confirming the functionality of different *Nin* transcripts in modulating strain-specific ethanol behavioral differences.

## Author Contributions

ERG and MFM conceived and designed the study. JLJ collected all tissue samples, designed PCR primers, and isolated nucleus accumbens RNA. ERG wrote manuscript, designed PCR primers, conducted all qRT-PCR, gel electrophoresis, DESeq2, DEXSeq, rMATs, and Regtools studies. MD assisted in RNAseq sample processing, quality control, and alignment. JTW assisted in PCR design and edited manuscript. MFM reviewed all data, edited manuscript and provided resources for conducting experiments.

## Acknowledgements

The data included in this study was generated at the Genomics Core facility at Virginia Commonwealth University. We would like to thank Kalyan Mallempati for cDNA library preparation and running the RNA-sequencing, and members of the Miles laboratory for helpful discussions in the preparation of this manuscript.

---

**Supplementary Table 1.**
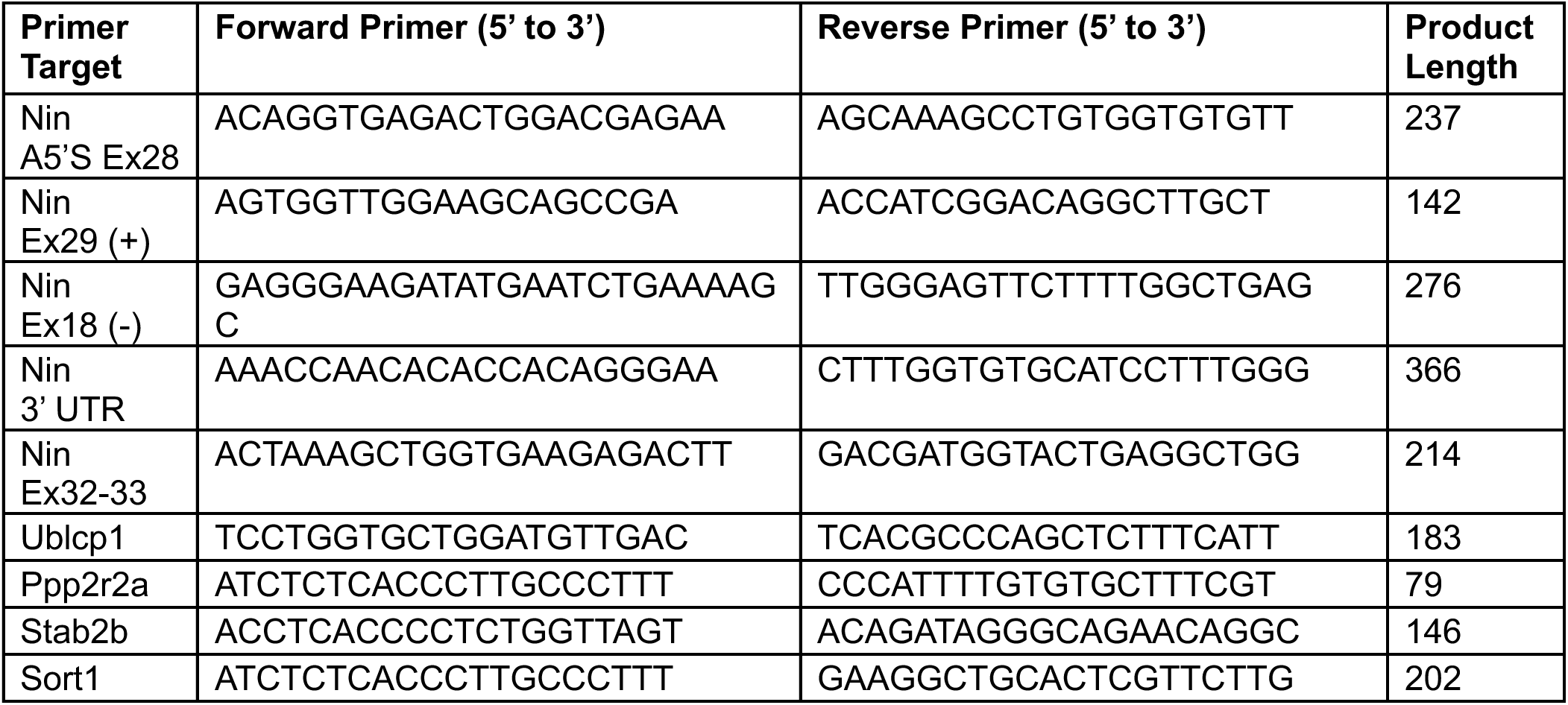
Primers used for quantitative reverse-transcriptase PCR (qRT-PCR). *Ublcp1, Ppp2r2a, and Sort1* were used as reference genes to calculate relative expression. *Stab2b* was used as a strain control.

**Supplementary Table 2.** RNA-sequencing information, quality control metrics, alignment percentages, and count generation analytics.

| Sample | RIN Value | Uniquely Mapped Alignment % | Assigned % |
| --- | --- | --- | --- |
| B14 N | 9 | 92.90% | 74.40% |
| B21 N | 9.1 | 91.60% | 74.90% |
| B24 N | 9.3 | 90.20% | 75.40% |
| B31 N | 9.1 | 90.90% | 74.60% |
| B32 N | 9.2 | 92.00% | 75.90% |
| D11 N | 9.3 | 92.50% | 75.00% |
| D13 N | 9.3 | 92.40% | 73.60% |
| D22 N | 9.1 | 92.40% | 74.40% |
| D32 N | 9.4 | 91.60% | 74.20% |
| D34 N | 9 | 91.20% | 74.60% |

**Supplementary Figure 1.**
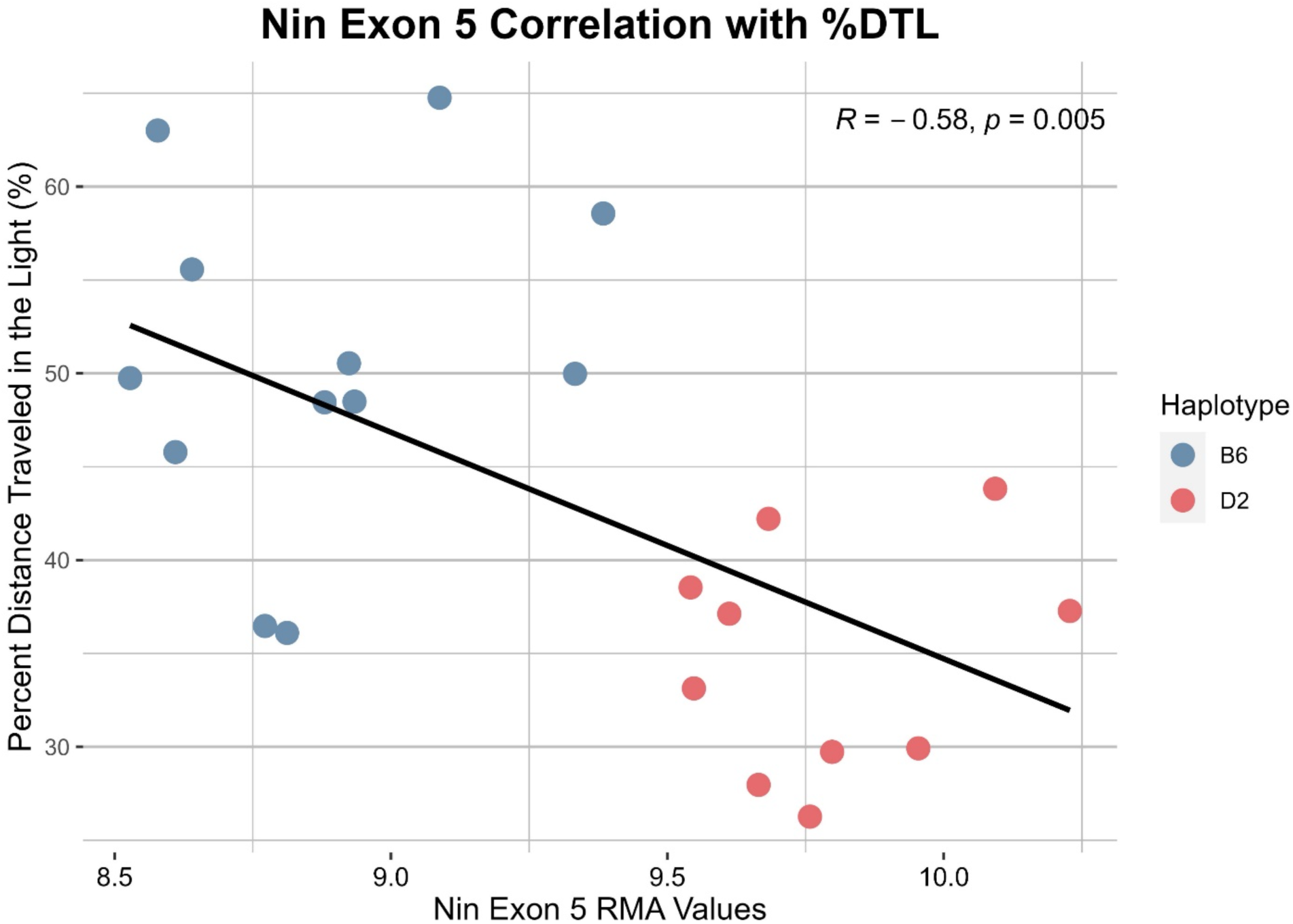
*Nin* Exon 5 Correlation with %DTL. Correlation of BXD strains and progenitors B6 and D2 with %DTL in response to 15-minute restraint stress and 1.8 g/kg EtOH (R = −0.58, p = 0.005 n = 22). BXD strains are highlighted by haplotype at *Nin* Exon 5.

## Other Supplemental Tables are available on request

**See** *Supp_Table_3_Nin_DEXSeq.xlxs*

**Supplementary Table 3. Ninein Differential Exon Utilization.** *Nin* exons that are significantly differentially utilized between B6 and D2 mice (FDR < 0.2).

**See**: *Supp_Table_4_Nin_rMATs.xlxs*

**Supplementary Table 4. Ninein Alternative Splicing.**

**See**: *Supp_Table_5_Nin_Junctions.xlxs*

**Supplementary Table 5. Ninein Splice Junction Analysis.**

**See**: *Supp_Table_6_Nin_SNPs.xlxs*

**Supplementary Table 5. Ninein Single Nucleotide Polymorphisms.**

